# Selective contexts favouring asymmetry: A theoretical and empirical study of pollen transfer in mirror-image flowers

**DOI:** 10.1101/2025.09.23.677969

**Authors:** Marco Saltini, Sam McCarren, Nicola Illing, Spencer CH Barrett, Bruce Anderson

## Abstract

Reciprocal herkogamy has evolved multiple times in flowering plants and is thought to enhance cross-pollination and reduce reproductive interference. Mirror-image flowers represent a form of reciprocal herkogamy in which plants have either right-or left-deflected styles (dimorphic enantiostyly) or produce both stylar orientations (monomorphic enantiostyly). The ecological conditions under which these two forms of enantiostyly originate and persist remain poorly understood. Here, we investigate how floral asymmetry affects pollen transfer dynamics and mating outcomes, and under which conditions enantiostyly provides a fitness advantage over straight-styled floral morphologies. We integrate field observations of dimorphic enantiostylous *Wachendorfia paniculata* with a mathematical model simulating pollen transfer in plants with straight-styled, monomorphic, and dimorphic enantiostylous flowers. The model incorporates pollinator pathways, pollen carryover, and floral display size, and was parameterized using our ecological data. Enantiostylous flowers received more outcrossed pollen than straight-styled flowers, with dimorphism outperforming monomorphism. However, enantiostyly increased stochasticity in pollen transfer due to variation in pollinator pathways. Intrafloral self-pollination was extremely low in enantiostylous flowers and did not differ between monomorphic and dimorphic forms. Dimorphic enantiostylous plants exhibited longer pollen carryover curves and exported more pollen to more mates potentially increasing siring success and mate diversity. Enantiostyly, particularly in dimorphic systems, can provide a selective advantage by improving pollen receipt and export components of fitness through enhanced outcrossing. This mating advantage may be reduced under conditions of pollen limitation or when autogamy in straight-styled flowers is low. Our findings clarify the functional significance of enantiostyly and offer a framework for understanding the evolution of floral asymmetry in angiosperms.

## 1 Introduction

Pollinators are key agents of floral diversification, driving the evolution of complex reproductive adaptations through their influence on pollen transfer and mating patterns (Harder and Barrett, 1996; Fenster et al., 2004; Johnson, 2025; Kay and Anderson, 2025). Diverse floral strategies function in a way that optimises cross-pollen transfer while, at the same time, minimizing the deleterious reproductive consequences associated with self-pollination. The reduction of mating costs is particularly important in animal-pollinated species with large floral displays that produce both pollen and ovules (Harder and Barrett, 1995). One of the great plant evolutionary dilemmas is that while increased display size can attract more pollinators and increase reproductive output, it also increases pollen movement between flowers of the same plant (geitonogamous self-pollination) (Klinkhamer and de Jong 1993). This potentially has negative consequences for pollen export and seed production, and is thought to have resulted in a multitude of evolutionary solutions.

Among the most complex floral adaptations are those involving stylar polymorphisms in which populations are subdivided into floral morphs that differ reciprocally in the positioning of reproductive organs, and outcrossing occurs primarily between morphs (disassortative mating). At least six different forms of reciprocal stylar polymorphism are reported in animal-pollinated angiosperm lineages, with heterostyly and enantiostyly as the most notable examples, since they are more frequent and widely dispersed among angiosperm families than the remaining polymorphisms (see Table 2 in Barrett, 2010). These two floral polymorphisms have been shown to promote effective cross-pollination, reduce pollen and ovule wastage and limit interference between reproductive organs (Webb and Lloyd, 1986; Kohn and Barrett, 1992; Barrett, 2002; Jesson and Barrett, 2005; Keller et al., 2014; Minnaar and Anderson, 2021; Simón-Porcar et al., 2015, 2024).

Since Darwin’s seminal studies (Darwin, 1877) heterostyly has received the most theoretical and empirical attention (reviewed in Ganders, 1979; Barrett, 1992; Simón-Porcar et al., 2024; Shang et al., 2025), but more recently enantiostyly has become a focus of significant research activity (Minnaar and Anderson, 2021; Barrett and Fairnie, 2024; Saltini et al., 2025; Robertson et al., 2025). In enantiostyly, styles are consistently deflected either to the left-or right-side of a flower (hereafter L- or R-flower or plant). This condition, also known as mirror-image flowers, is reported from at least 10 angiosperm families particularly associated with heteranthery (Jesson and Barrett, 2003). This floral polymorphism exists in two major forms: monomorphic enantiostyly, where individual plants produce both L- and R-styled flowers, and dimorphic enantiostyly, where plants are fixed for a single stylar orientation and populations are polymorphic for L- and R-individuals (see Fig. 5 in Barrett et al., 2000). The vast majority of enantiostylous species are monomorphic, and dimorphic enantiostyly is only known from a handful of species, including members of *Wachendorfia, Barberetta* (Haemodoraceae) and *Heteranthera* (Pontederiaceae) (Ornduff and Dulberger, 1978; Jesson and Barrett, 2002c; Johnson et al., 2023). While monomorphic enantiostyly is often associated with and pollination by pollen-collecting bees (Jesson and Barrett, 2003), most dimorphic species (all species from the genera *Wachendorfia* and *Barberetta*) produce nectar (Barrett and Fairnie, 2024).

The overall rarity of dimorphic enantiostyly compared to monomorphic enatiostyly raises intriguing questions about the conditions necessary for its evolution and persistence. Phylogenetic analyses and modelling approaches indicate that monomorphic enantiostyly most likely arises from straight-styled ancestors and that dimorphism evolved secondarily (Jesson et al., 2003a; Saltini et al., 2025). Transitions may be constrained by developmental genetics, e.g. limited positional cues for left-right organ placement, or by weak selection for strict reciprocity, especially in species with small floral displays (Barrett et al., 2000; Jesson et al., 2003b). A further complication concerning the relative frequency of enantiostylous conditions is that monomorphic enantiostyly, by producing flowers of both stylar orientations on the same plant, has been shown experimentally to incur higher rates of geitonogamy than dimorphic enantiostyly (Barrett et al., 2000; Jesson and Barrett, 2002b), which can potentially lead to inbreeding depression and fitness losses through pollen discounting, i.e. self-pollination reducing the amount of pollen available for outcrossing (Lloyd, 1992; Harder and Wilson, 1998). The reproductive organs within enantiostylous flowers are spatially separated (herkogamy) and therefore levels of within-flower self-pollination may often be reduced compared to straight-styled plants (Webb and Lloyd, 1986; Jesson and Barrett, 2002b). Indeed, this reduction in within-flower self-pollination resulting in autogamy may have been the initial advantage favouring the evolution of monomorphic enantiostyly, with selection for fixed handedness in dimorphic enantiostyly emerging later as adaptations that reduce geitonogamy and enhance pollen export and outcrossing (Barrett et al., 2000; Jesson and Barrett, 2002b). Dimorphic enantiostyly should promote mating between L- and R-individuals (and vice versa) in species where pollinators deposit pollen on specific body regions (Minnaar and Anderson, 2021; Johnson et al., 2023). Thus, sequential visits to the same morph might only deposit small quantities of pollen, resulting in increased pollen carryover and consequently larger genetic neighbourhoods (Karron et al., 2021), which likely reduces self-pollination between flowers of the same plant.

These functional hypotheses and their role in the evolution of enantiostyly have been investigated in mathematical models (Jesson et al., 2003a; Jesson and Barrett 2005; Saltini et al., 2025). Pollen transfer between morphs might lead to increased fitness but this is dependent on several factors including floral display size, pollinator behaviour, inbreeding depression, and the efficiency of pollen pickup and deposition by insects. Theoretical models have shown that enantiostyly can reduce geitonogamy relative to straight-styled flowers and that dimorphism is more likely to evolve under high pollinator visitation, strong inbreeding depression, and an abundance of abiotic resources (Jesson et al., 2003a; Saltini et al., 2025). However, the conditions under which monomorphic or dimorphic forms might be favoured in particular ecological circumstances remain poorly understood. One possibility that has received little attention is that monomorphic enantiostyly might arise not as a mechanism that reduces geitonogamy compared to straight-styled ancestors, but rather because it limits within-flower selfing through increased herkogamy. This hypothesis could be particularly relevant in species with few flowers open at a time. In such cases, geitonogamous selfing is already minimized by small floral displays, so within-flower selfing becomes the dominant source of inbreeding. This might help explain the evolution of monomorphic enantiostyly in taxa like *Cyanella alba* (Dulberger and Ornduff, 1980; Robertson et al., 2025) or *Streptocarpus* (Harrison et al., 1999) where small floral displays reduce geitonogamous selfing (see Table 2 in Karron et al., 2003). Under this hypothesis, monomorphic enantiostyly could evolve even in the context of low pollinator service and limited cross-pollination, whereas subsequent shifts to dimorphic enantiostyly would be favored once larger floral displays and higher pollinator visitation make geitonogamy more costly. To illuminate how enantiostylous traits evolve and persist, it is necessary to understand the fitness consequences of pollen dispersal. Plant fitness depends on both pollen receipt and pollen export, both of which are influenced by pollinator behaviour, including the number of flowers visited per plant, the fraction of pollen deposited on stigmas, and the spatial segregation of pollen on pollinator bodies (Iwasa et al., 1995; Harder and Barrett, 1996; Harder et al., 2000).

Here, we extend the mathematical model originally introduced by Jesson and Barrett (2005) to incorporate pollen import and export in a focal plant exhibiting either dimorphic enantiostyly, monomorphic enantiostyly, or a straight-style condition. In contrast to the original model, we account not only for geitonogamous pollen transfer but for the full spectrum of intra-specific pollen movement, i.e., pollen carried from one plant to subsequent plants, including losses due to deposition on intervening plants between the donor and the focal recipient. In addition, we incorporate intrafloral self-pollination. We combine our mathematical models with field observations of *Wachendorfia paniculata*, a dimorphic enantiostylous species, to investigate the ecological and selective conditions under which monomorphic and dimorphic enantiostylous traits might confer fitness advantages.

The main aim of our study is to explore how 1) within-flower self-pollination, 2) pollinator behaviour, and 3) floral display size influence fitness through pollen export and import in straight-styled, monomorphic, and dimorphic floral forms. We expect that enantiostyly leads to increased pollen carryover, particularly for dimorphic forms. Further, we predict reduced within-flower self-pollination in both monomorphic and dimorphic enantiostylous plants compared to straight-styled plants. Both forms of enantiostyly are likely to outperform straight-styled plants when floral displays are small or when pollinators visit few flowers and deposit little pollen. However, the advantage of the monomorphic condition might decline when pollinators visit more flowers per inflorescence or deposit a larger fraction of pollen, thereby increasing the relative cost of geitonogamous self-pollination. On the other hand, we expect the dimorphic form which is known to suffer less geitonogamous self-pollination to maintain a reproductive advantage across a wider range of conditions. By linking ecological data with evolutionary theory, we seek to clarify the pathways and constraints underlying the evolution of floral asymmetry in angiosperms.

## 2 Methods

### 2.1 Field observations

We conducted field studies of a population of *W. paniculata*in 2024 at Stellenbosch Mountain (−33.9406, 18.8864) in the Western Cape Province, South Africa. The focal site had burned recently and there was a high density of flowering *W. paniculata* plants abundant over the entire area of approximately 200 hectares. We estimated that the population size at our study site was approximately 2000 individuals. We measured density (P) by counting the number of *W. paniculata* plants in 15 randomly chosen 10 *m*^2^ plots and also recorded the style morph ratio (M) and average daily display size (D) for 100 randomly chosen individuals in the population. We estimated pollen production (*α*) by randomly collecting the anthers from 30 newly opened, unvisited flowers which were stored in ethanol for later counting the amount of pollen produced for each flower using a haemocytometer under a compound microscope at 40× magnification (Kearns and Inouye, 1993). To determine features of pollinator visitation, we recorded the identity of pollinators, the number of flowers a pollinator visited per plant (*v*) and the approximate distance if flew to the next plant (*d*). These data were then compared between the primary pollinator species honeybees (*Apis mellifera*) and carpenter bees (*Xylocopa caffra*) (Figure 1) using Mann-Whitney U tests in R (R Core Team, 2020). To investigate pollen movement, we tracked pollen deposition from focal donor flowers to a sequence of recipient flowers. A total of 18 focal plants and 80 flowers were covered with pollinator exclusion bags in the evening before the experiment to prevent any pollen deposition or removal before the start of the experiment. Once anthers had fully dehisced in the morning, we labelled all flowers on a focal plant with quantum dots (q-dots) following methods described by Minnaar and Anderson (2019). Q-dots are nanoparticles that fluoresce under UV light excitation. After labelling all flowers on a focal plant, we removed the pollinator exclusion bag and allowed pollinator visitation. After a pollinator visited one of the focal flowers, it was gently ushered away from the focal plant to prevent it from visiting more than one labelled flower. We observed the pollinator’s subsequent pathway in the population up to the sixth recipient flower after the donor flower or until we lost sight of it (honeybees *n*= 78, carpenter bees *n*= 2). The recipients flowers were not covered, which means that they often had received previous visits, however the focal pollinator carrying labelled pollen only visited them once. We then collected and recorded the morph of both the donor flower, as well as the recipients and their stigmas, which were placed on a slide and secured under a cover slip using sticky tape for later analysis. This was repeated for all flowers on a focal plant before moving on to the next focal plant. In the laboratory, we counted the total number of pollen grains on stigmas using a compound microscope at 40× magnification. We also identified and counted q-dot labelled pollen grains on stigmas using the same compound microscope and a handheld UV torch. We analysed pollen movement in R (R Core Team, 2020). Since donor flowers received only one visit, the total pollen on their stigmas represented single visit deposition (*R*_0_) and the relative amount of labelled pollen represented within-flower self-pollination (*λ*_*E*_). We fitted a generalized linear model from the package glmmTMB (Brooks et al., 2017) with Poisson error structure and correction for zero-inflation to quantify single visit deposition in response to pollinator species. To estimate pollen deposition curves, we analysed the amount of q-dot labelled pollen transferred from the donor flower to the stigmas that were sequentially visited by the pollinator (*R*). We further repeated this model with an interaction term between visitation sequence and whether the recipient flower was the same or a different morph from the labelled donor flower. Additionally, to inform the parameter estimates used in the mathematical model we performed further analyses of data collected by Minnaar and Anderson (2021) in their quantum dot study of *W. paniculata*. From their supplementary table we extracted the total number of pollen grains picked up from a flower by each pollinator (*R*), as well as the contribution to this number by the upper anther (*R*_*u*_), the lower anther on the same side as the stigma 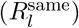 and the lower anther on the other side 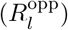.

**Figure 1.**
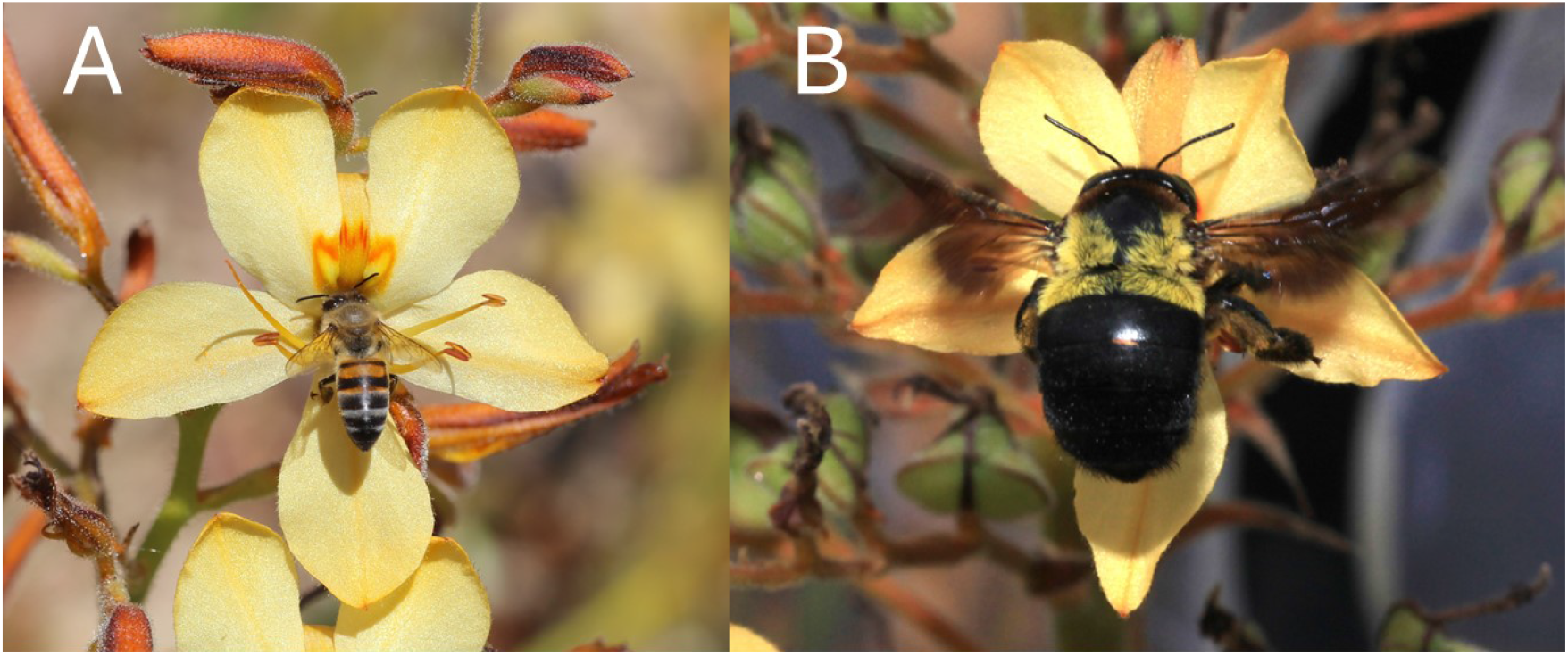
*Wachendorfia paniculata* (A) visited by honeybee (*Apis mellifera*) and (B) visited by carpenter bee (*Xylocopa caffra*)

### 2.2 The model

Our model, in its general formulation, tracks the pollen exported and imported by a focal plant in a visitation sequence by a generic pollinator. We investigated three different general scenarios: i) the focal plant has dimorphic enantiostyly as in *W. paniculata*, ii) the plant has monomorphic enantiostyly, iii) the plant has both a straight-style and straight pollinating anthers. In the dimorphic case, for simplicity, we assume that the focal plant has a left-deflected style, our results would be equally applicable and give the same results if the plant was right-handed.

#### 2.2.1 Model assumptions and parameters

In our model, each time a pollinator visits a flower it collects Π pollen grains. We assume that exactly *R* of those grains survive transport to the next flower in the visitation sequence. For simplicity, *R* is taken to be independent of the distance between successive flowers and, hereafter, we refer to such quantity as pollen collected by a pollinator. In the enantiostylous case, the incoming pollen is stored in three mutually isolated compartments on the pollinator’s body, depending on the relative position of the pollinating anthers and stigma of a flower (see Minnaar and Anderson, 2021): (i) the top part of the side of the body opposite to the stigma, (ii) the bottom part of the side of the body opposite to the stigma, and (iii) the bottom part of the same body side as the stigma. Pollen originating from different donor flowers is assumed to mix thoroughly *within* each compartment, so a recipient flower encounters a well-mixed sample of all previously visited donors. For straight-styled flowers, we assume a single compartment where pollen from all pollinating anther is stored.

During a visit a fixed fraction *ρ* of the pollen present on the body side that contacts the stigma is deposited. Grains on the opposite side never reach the stigma (e.g., an L-flower cannot receive pollen from the R-side of a pollinator). Intraflower self-pollonation is represented by the parameter *λ*, the proportion of pollen that moves directly from a flower’s own anthers to its stigma. Finally, we assume that a pollinator visits exactly *v* flowers on each plant before departing. Therefore, it is possible to define the function *q* as the fraction of pollen on a pollinator’s body that is deposited on a plant after visitation by using a geometric sum: a fraction *ρ* of that load is deposited on the first flower of plant *N*, a fraction *ρ*(1 − *ρ*) on the second, *ρ*(1 − *ρ*)^2^ on the third, and so on. Therefore,

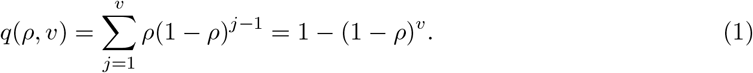

We use data obtained from our field observations to numerically solve our models in the case in which the visiting pollinator was *Apis mellifera*. Relevant parameter values used in our models are listed in Table 1 and are based on our field data collection (see Section 2.2).

**Table 1.**
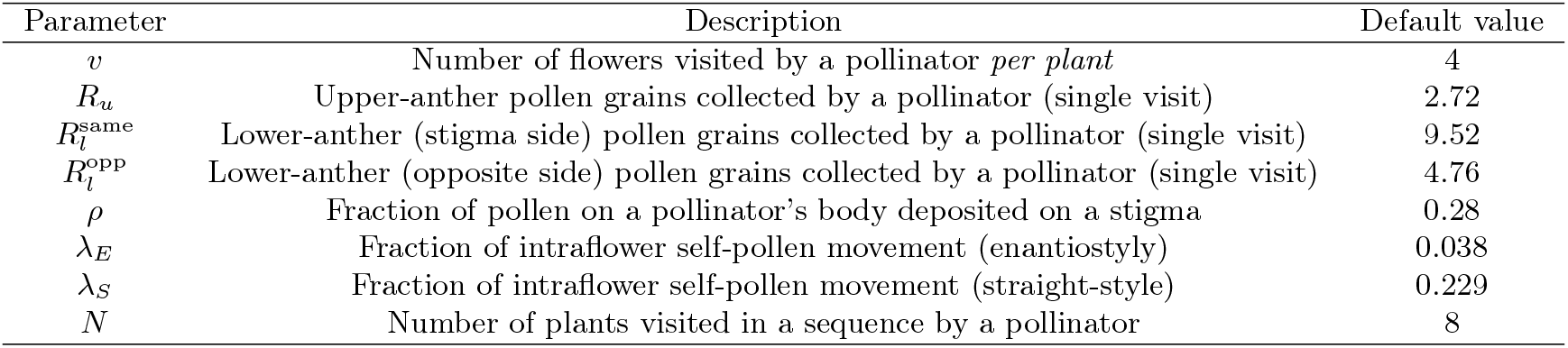
Default model parameters.

#### 2.2.2 Geitonogamous pollen transfer

As in our model we assume that pollen remains compartmentalised on a pollinator’s body and grains are deposited only from the side of the body that contacts the stigma, in enantiostylous plants geitonogamous transfer involves exclusively the lower anther on that same side. Building on the expressions of Jesson & Barrett (2025) and adding the intraflower self-pollination component *Rvλ*, we obtain the geitonogamous pollen transfer for both the dimorphic case 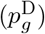 and the straight-styled case 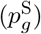 as

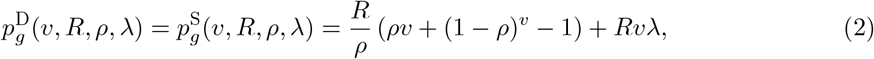

In the monomorphic case, instead, the amount of geitonogamous pollen transferred 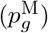 is

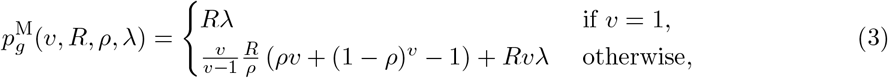

where the *v* = 1 case is treated separately because the term for *v ?*= 1 is undefined when *v* = 1. These formulas give the expected proportion of geitonogamous pollen transferred under each stylar condition, incorporating contribution for both inter-floral and intra-floral self-pollination.

#### 2.2.3 Pollen imported from other plants

We compute the pollen imported by the *N* -th plant visited in a pollination sequence.

For plants with straight-styled flowers, the imported load equals the pollen picked up from each of the plants *j* = 1, …, *N* − 1 visited before *N*, minus the fraction subsequently removed on the later plants *k* = *j* + 1, …, *N* − 1. Therefore, we can show that the pollen received by plant *N* is

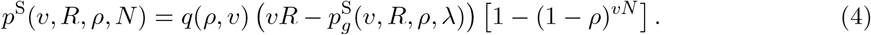

In the case of dimorphic enantiostyly, we distinguish between pollen transferred from *n* plants of the same symmetry, i.e., *p*^same^, and those *N* − *n* plants of the opposite symmetry, i.e., *p*^opp^ relative to the target L-plant. Therefore, pollen imported is parametrized as

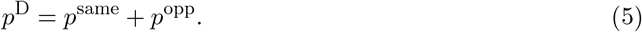

When the source is an L-flower, only the lower anther contributes to pollen transfer. When a pollinator visits a L-plant, it collects a total of *vR* pollen grains, with 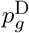 grains that are deposited on the stigma of the same plant. The remaining 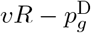 grains from plant are available to the other *n* − 1 L-plants. After each visited flower on those plants, a fraction (1 − *ρ*) is lost through deposition on stigmas. Hence the amount of pollen carried immediately before the pollinator lands on the first flower of plant *N* is

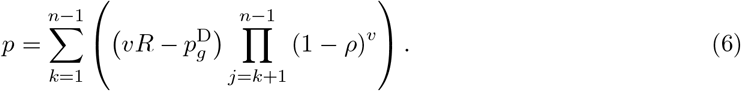

By solving the product first, and the sum later, we can compute the total pollen received by plant *N* from other L-plants:

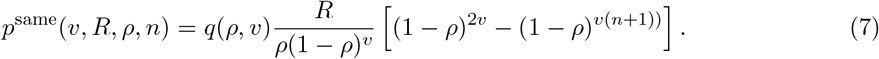

When pollen originates from R-plants, the amount deposited on the focal L-plant depends on the precise visitation pathway *t*(*n, N*) of the pollinator. For example, if a pollinator visits four flowers in the order L, L, R, L, all the pollen it collected on the third (R) flower reaches the final L flower, and the latter receives *Rρ* grains. Conversely, with the sequence R, L, L, L, the grains collected on the first R flower are progressively depleted on the second and third flowers, so the last L flower receives only *Rρ*(1 − *ρ*)^2^ grains. In general, the pollen received from opposite-symmetry plants by the focal plant is

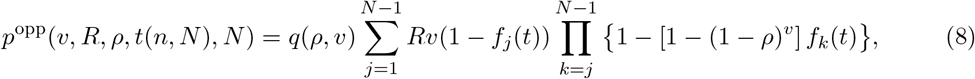

where

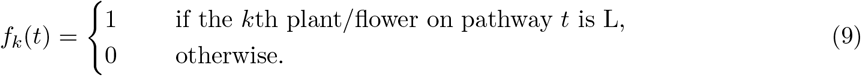

The summation captures the pollen collected on each R-plant (terms with (1 − *f*_*j*_)), whereas the product accounts for losses on subsequent L-flowers along the pathway. We are interested in the average of *p*^opp^ over all (*N* − 1)! possible pathways:

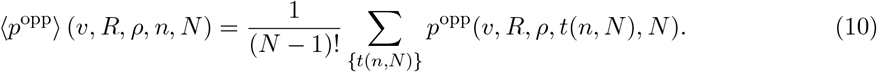

Therefore, the average total number of pollen grains received by our focal L-plant, is

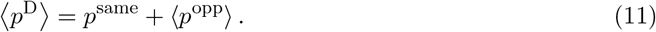

To calculate the pollen received by a monomorphic plant, we treat a plant with *v* flowers as if it was composed of *v* single-flowered plants. In this case, a pollinator follows *N* − 1 different pathways (*t*_1_, *t*_2_, …, *t*_*N*−1_), one per visited plant before visiting the focal plant. Using the dimorphic results under this decomposition, we find

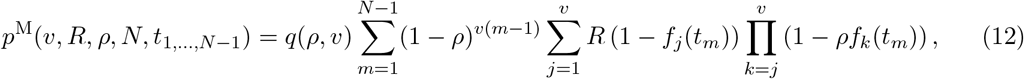

and

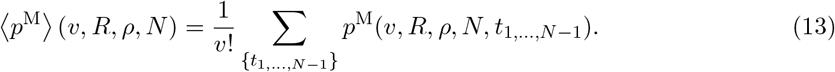

##### Fraction of pollen imported

To quantify how efficiently each floral architecture imports pollen, we define the proportion of pollen removed from earlier plants that ultimately reaches the stigmas of a focal plant during a single pollinator visitation sequence compared to the total pollen received as:

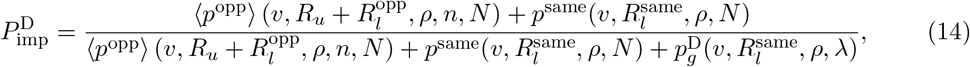

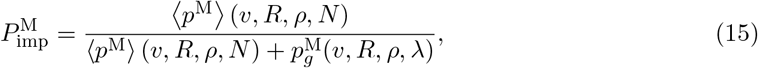

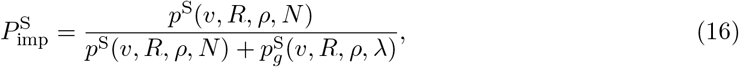

for dimorphic, monomorphic, and straight-styled cases, respectively. Note that in the dimorphic case, we retained the separate contributions of the upper anther (*R*_*u*_) and the lower anthers on the opposite 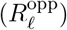 and same 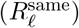 sides. In the monomorphic and straight-styled case, instead, we used the total pollen collected by a pollinator 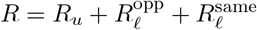 .

#### 2.2.4 Pollen exported to other plants

We now compute the average number of pollen grains *exported* from a focal plant to the subsequent plants labelled *k* = 1, 2, … in a pollinator’s visitation sequence.

In straight-styled flowers a pollinator removes 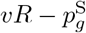 grains from the focal plant. A fraction *h* = 1 − *q*(*ρ, v*(*k* − 1)) = (1 − *ρ*)^*v*(*k*−1)^ is lost on the stigmas of plants 1, …, *k* − 1. Consequently, the *k*-th plant receives

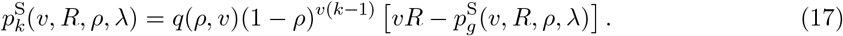

For dimorphic enantiostyly the exported load also depends on the pollinator’s pathway. Averaging over the 2^*k*^ possible sequences of L- and R-handed plants among the *k* recipients, we obtain

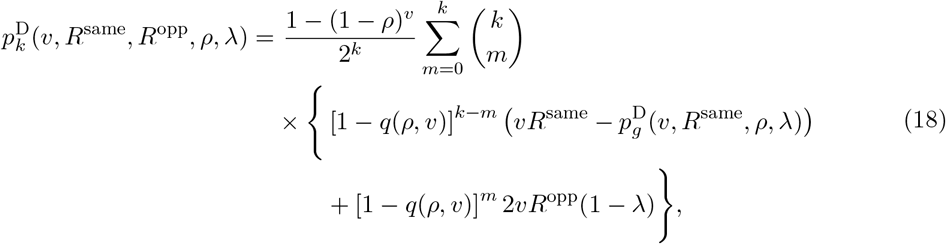

where the first term in braces accounts for losses on plants of the same symmetry and the second for those of the opposite symmetry. The factor 2 reflects the two pollinating anthers that supplied the pollen, whereas losses on the focal plant itself occur only via intra-floral self-pollination, represented by *λ*.

Because a monomorphic plant functions like a straight-styled plant with *v* equivalent flowers, the amount exported to the *k*-th recipient is

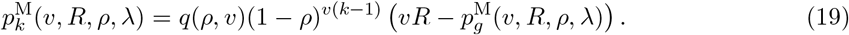

##### Fraction of pollen exported

To quantify how efficiently each floral architecture exports pollen, we define the proportion of pollen exported by the focal plant to the subsequent plant compared to the pollen loss in geitonogamy as:

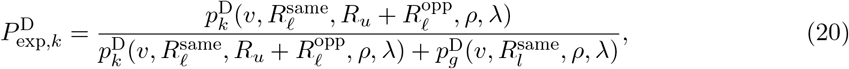

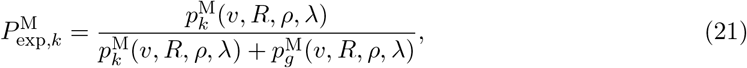

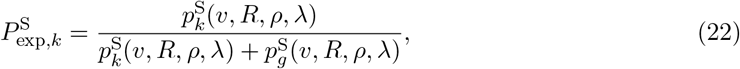

for dimorphic, monomorphic, and straight-styled cases, respectively. As before, note that in the dimorphic case, we retained the separate contributions of the upper anther (*R*_*u*_) and the lower anthers on the opposite 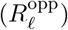 and same 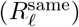 sides. In the monomorphic and straight-styled case, instead, we used the total pollen collected by a pollinator 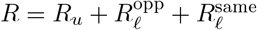 .

## 3 Results

### 3.1 Field observations

The density of *W. paniculata* at our study site was ≈ 2.3 ± 0.9 plants/*m*^2^ and the average display size was ≈ 12.6 ± 8.8 open flowers/plant. The observed morph ratio was 54L-plants:46R-plants (*n* = 100) and morphs appeared to be randomly distributed since we did not detect any evidence of significant spatial clumping.

Pollen production per flower was *α* ≈ 6410 ± 3870 pollen grains. We observed both honeybees (*Apis mellifera*), which were by far the most frequent visitors, as well as carpenter bees (*Xylocopa caffra*) visiting *W. paniculata* flowers (Figure 1). Carpenter bees visited *v*_*C*_ ≈ 2.8 ± 2.0 flowers/plant which was significantly less than honeybees which visited *v*_*H*_ ≈ 3.8 ± 3.1 flowers/plant (*W* = 15864, *p <* 0.001). Additionally, carpenter bees flew longer distances of approximately *d*_*C*_ ≈ 1.5 ± 2.1 m between plants in comparison to honeybees which flew approximately *d*_*H*_ ≈ 0.7 ± 0.5 m (*W* = 20677, *p <* 0.001).

Single visit pollen deposition (including zeros) from honeybees was *R*_0,*H*_ ≈ 1.5 ± 5.3 pollen grains with a within-flower self-pollination of *λ* ≈ 3.79 %, whereas carpenter bees deposited an average of *R*_0,*C*_ ≈ 2.0 ± 2.8 pollen grains (*n* = 2). There were insufficient visits from carpenter bees for a statistical comparison of single visit deposition or to establish a pollen deposition curve. However, for honeybee visits, there was a significant effect of visitation sequence on pollen deposition (*χ*^2^ = 5.19, df = 264, *p* = 0.023). As the number of flowers visited after the donor flower increased, the amount of pollen transferred decreased (Fig. 2). The curve fitted for honeybee pollen deposition was

**Figure 2.**
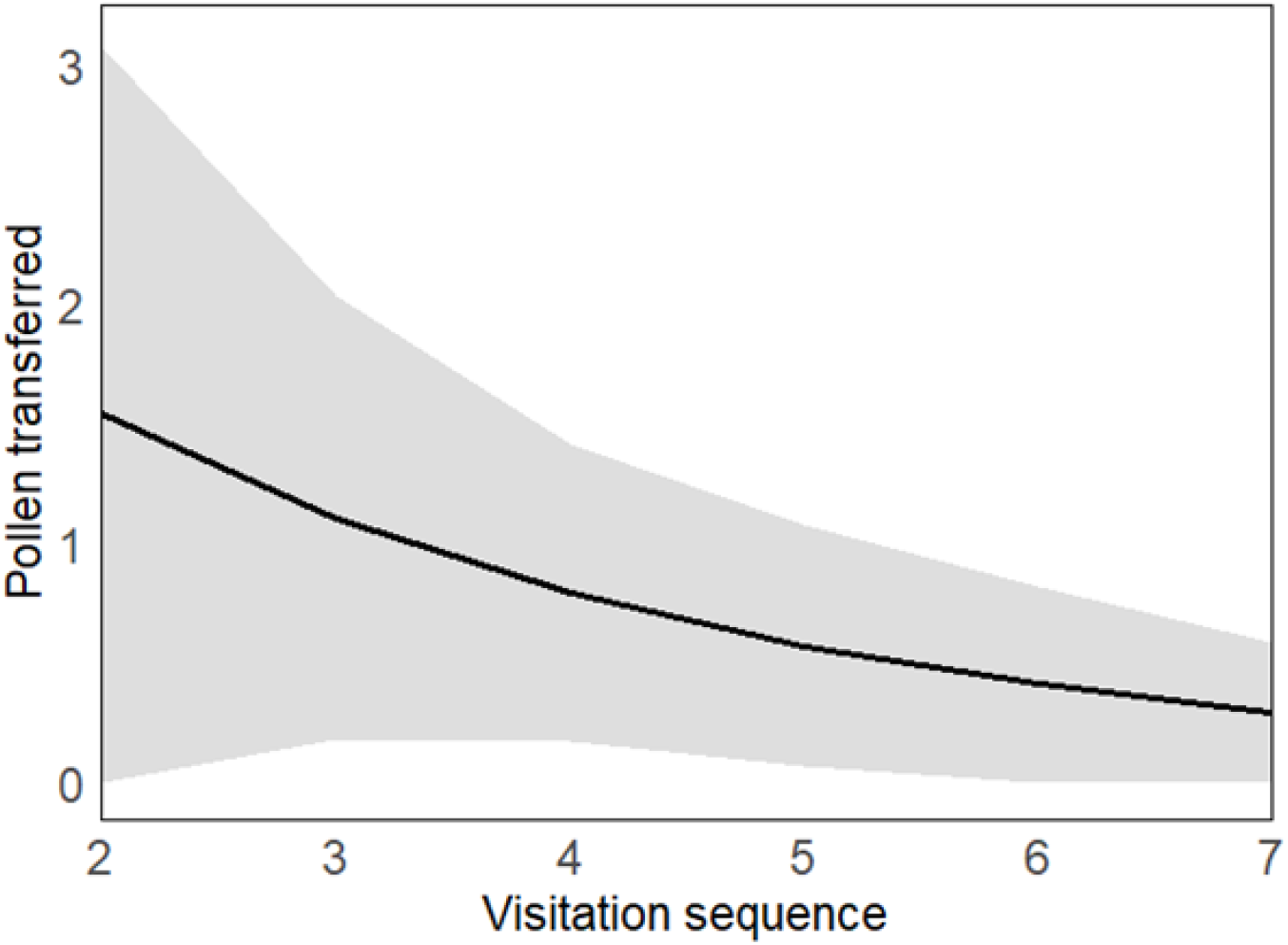
Field study of pollen transferred (*p*_*k*_) from a focal *Wachendorfia paniculata* flower to recipient flowers in relation to the sequence (*k*) they were visited including visits within and between morphs.

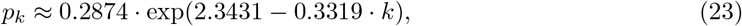

where *k* = 1, 2, … is the position of a flower in the sequence. Note that our fitted curve as in Figure 2 is qualitatively similar to the theoretically derived curve for pollen export (Figure 4). The model including an interaction between visitation sequence and within or between morph visits showed a similar effect of sequence on pollen deposition (*χ*^2^ = 7.66,df = 262, *p* = 0.006), whereas morph alone was not significant (*χ*^2^ = 1.13, *p* = 0.287). The interaction of the terms, while only being close to significance (*χ*^2^ = 2.93, *p* = 0.087) still indicated that the pollen deposition curve likely decreased faster for visits within a stylar morph and more gradually for visits between morphs. Our analysis of the data collected by Minnaar and Anderson (2021) indicated that carpenter bees picked up *R*_*C*_ ≈ 66 pollen grains/visit whereas honeybees picked up *R*_*H*_ ≈ 17 pollen grains/visit. The contribution of the upper anther was *R*_*u*_*C* ≈ 68 % in carpenter bees and *R*_*u*_*H* ≈ 8 % in honeybees. The anther on the same side as the style contributed 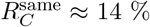 for carpenter bees and 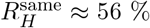 for honeybees, while the lower anther on the other side contributed 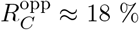 for carpenter bees and 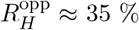 for honeybees.

### 3.2 The model

#### 3.2.1 Dimorphic enantiostyly promotes greater pollen import and export

Dimorphic enantiostyly resulted in greater import of outcross pollen than monomorphic enantiostyly which, in turn, outperformed the straight-styled condition (Figure 3). The illustrated values were calculated at the end of a visitation sequence involving 10 plants. Figure 3A revealed that both monomorphic and dimorphic enantiostyly were more effective at promoting cross-pollination when pollinators visited fewer flowers per plant, and increased flower visitation elevated levels of geitonogamous self-pollination (see also Jesson & Barrett, 2005). In contrast, for straight-styled plants, there existed an optimal number of flowers visited (for parameter values as in Table 1) that maximized pollen import. Figure 3B indicated that increasing the fraction of pollen deposition (*ρ*) initially boosted pollen receipt, but beyond a certain point, it began to decline. This response was because higher deposition led to more pollen being left on the first plant visited, thereby reducing transfer to subsequent plants. Importantly, under high values of both *v* and *ρ*, the outcomes for monomorphic and straight-styled plants were comparable, suggesting that the key advantage of dimorphic enantiostyly lies in its ability to strongly reduce geitonogamy due to reduced left to left and right to right pollen transfers, and thereby enhancing outcross pollen receipt.

**Figure 3.**
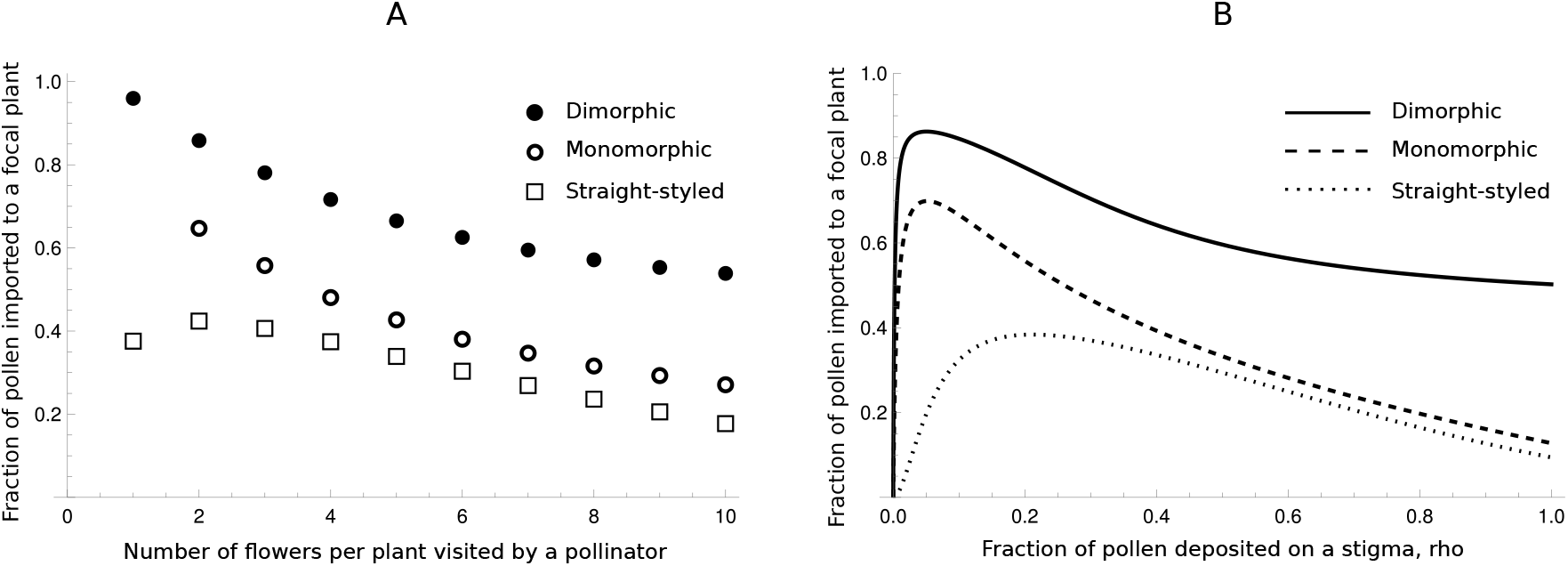
Model results illustrating the average fraction of outcrossed pollen received by a focal plant calculated over all possible pollinator pathway as a function of: (A) the number of flowers visited per plant by the pollinator (*v*), and (B) the fraction of pollen on the body of a pollinator deposited on the stigma of the focal plant (*ρ*). Solid lines and filled circles represent dimorphic enantiostyly, dashed lines and empty circles represent monomorphic enantiostyly, and dotted lines and empty squares represent straight-styled species.

Pollen import for dimorphic enantiostyly was highly sensitive to the specific pathway taken by a pollinator (see Figure S2). Therefore, although dimorphic enantiostyly tended to outperform the other stylar conditions on average, it may not always be the most effective floral strategy for increasing pollen import from single pollinator flights, especially when pollinator visits are rare.

Dimorphic enantiostyly also promoted pollen export across a visitation sequence (Figure 4) and consistently outperformed monomorphic enantiostyly which, in turn, outperformed the straight-styled condition. Specifically, the model predicted that in the dimorphic case, the proportion of exported pollen (relative to geitonogamously transferred pollen) becomes negligible after 10 plant visits, compared to only 3 plant visits for both the monomorphic and straight-styled scenarios.

**Figure 4.**
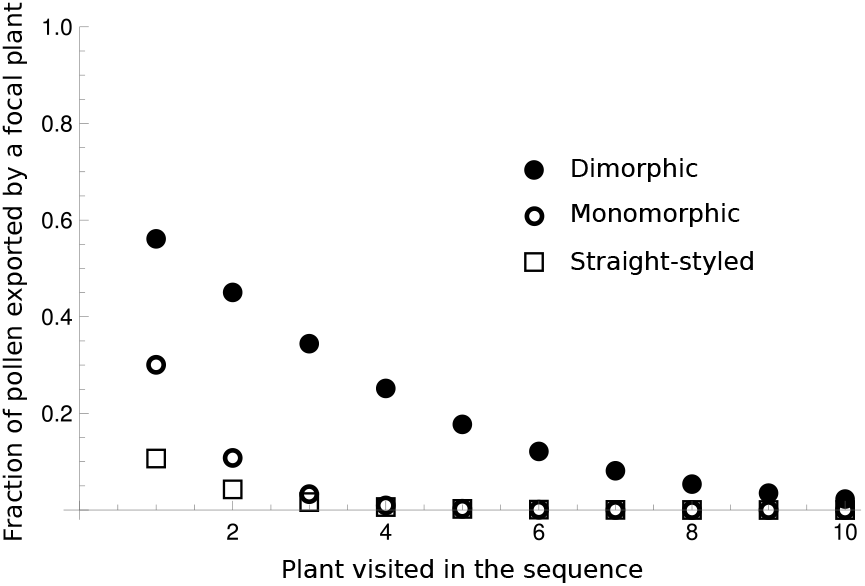
Model results of the average fraction of outcrossed pollen exported by a focal plant to the 1st through 10th plant in the visitation sequence of a pollinator. Filled circles represent dimorphic enantiostyly, empty circles represent monomorphic enantiostyly, and empty squares represent straight-styled species.

#### 3.2.2. High levels of intrafloral self-pollination reduce both import and export of pollen

Figure 5 illustrates the fraction of pollen imported to and exported by a straight-styled focal plant as a function of the degree of intrafloral self-pollination *λ*_*S*_, compared to both dimorphic and monomorphic enantiostyly. As predicted, both pollen import and export decreased with increasing intrafloral self-pollination in the straight-styled case. When this form of self-pollination was very low, straight-styled plants imported more pollen than monomorphic enantiostyly, although values remained lower than in the dimorphic case (Figure 5A). Conversely, for the parameter values considered, pollen export in straight-styled plants was always lower than in enantiostylous plants, even when intrafloral self-pollination was absent (Figure 5B). However, in the case of carpenter bees, there exists a value of *λ*_*S*_ at which pollen export in the straight-styled condition exceeds that of monomorphic enantiostyly (see Figure S1). Overall, dimorphic enantiostyly consistently outperforms the straight-styled condition, even when intrafloral self-pollination was absent in the latter condition.

**Figure 5.**
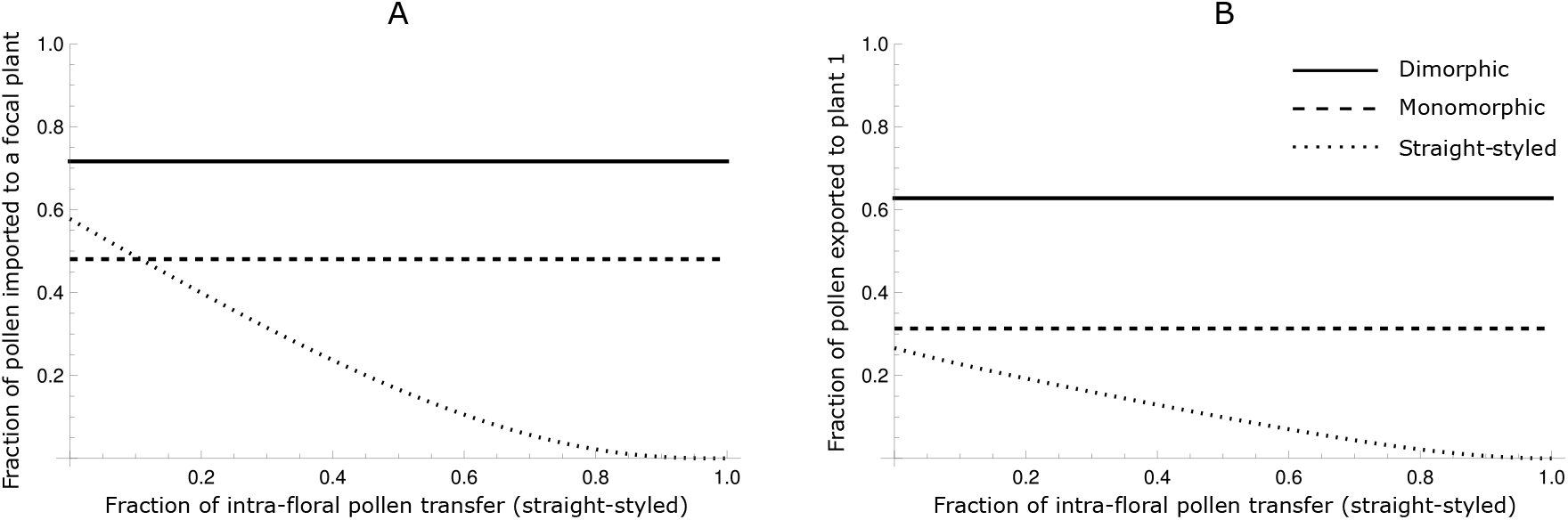
Model results of the average fraction of outcrossed pollen (A) received by a focal plant and (B) exported to the subsequent plant in the pathway of a pollinator, as functions of the fraction of intrafloral pollen transfer in straight-styled species (*λ*_*S*_). Solid lines represent dimorphic enantiostyly, dashed lines represent monomorphic enantiostyly, and dotted lines represent straight-styled species.

## 4 Discussion

Our results demonstrate that enantiostyly can increase pollen carryover and both our field data and model indicated that dimorphic enantiostylous plants had a much longer pollen carryover curve than either monomorphic or straight-styled plants (Figures 4 and 2). Pollen was retained on pollinators and successfully exported over more successive visits to flowers, increasing both the number of seeds sired and potential mate diversity. The consistently higher pollen export in dimorphic systems compared to the other stylar conditions likely has strong fitness consequences under conditions of high visitation rates and weak pollen limitation, where pollen competition among donor morphs is likely to become a key driver of fitness (Harder et al., 2000). Interestingly, Lloyd and Webb (1992) had suggested that heterostylous polymorphisms may increase pollen carryover in populations, but without support. Our study provides empirical and theoretical evidence in line with this suggestion and raises the possibility that increasing pollen carryover and the resulting fitness benefits may be a fundamental feature of stylar polymorphisms.

In contrast, straight-styled and monomorphic enantiostylous plants exhibited shorter carryover curves, with most pollen deposited within the first few flowers visited. Much of this pollen was transferred between flowers on a plant resulting in limited pollen export to other plants. Nevertheless, our model indicates that monomorphic enantiostylous plants export more pollen than the straight-styled condition, offering a modest advantage, especially under conditions of moderate to high self-pollination. These patterns are consistent with those of (Jesson and Barrett, 2005), who found that estimated geitonogamy, based on genetic markers, was highest in straight-styled individuals, intermediate in monomorphic individuals, and lowest for the dimorphic condition.

However, our results also indicated that pollen transfer in enantiostylous systems may be more variable, owing to the stochastic nature of pollinator trajectories (Fig. S2). In our model, the efficiency of pollen delivery in both monomorphic and dimorphic enantiostylous species was sensitive to the sequence of floral visitation. In contrast, straight-styled plants in our models were not affected by variation in pollinator movement, because all flowers presented their stigmas and anthers in roughly the same direction. This uniform flower morphology makes pollen transfer in straight-styled plants less variable, albeit less efficient in terms of outcrossing. Increased stochasticity in enantiostylous systems could pose a risk when pollinator activity is low or when visitation rates are erratic, potentially reducing reproductive reliability, especially in small populations (Harder and Barrett, 1996; Ashman et al., 2004).

Despite the potential vulnerability to stochastic pollination enantiostylous species have evolved floral morphologies (e.g. well-developed herkogamy) that limit within-flower self-pollination. Our field observations revealed that within-flower self-pollination was extremely low in *W. paniculata*, which is expected to be the case in all enantiostylous plants regardless of whether they possess, monomorphic or dimorphic systems. Our results are consistent with (Jesson and Barrett, 2005) who found that rates of within-flower self-pollination in enantiostylous flowers were substantially lower than in comparable straight-styled species. Our findings are relevant to a broader literature on pollen deposition in angiosperms: studies in non-enantiostylous species report within-flower pollen deposition rates ranging from 8.5% to 45% (e.g. Karron et al., 2003; Hou and Zhao, 2023; Chen and Pannell, 2024); in contrast we found values that were an order of magnitude lower in our focal enantiostylous species.

Significantly, in our models the advantage of enantiostyly increased under high levels of within-flower self-pollination in straight-styled species (Fig. 5). When self-pollen deposition was low in straight-styled plants, the relative benefit of monomorphic enantiostyly became negligible. However, as within-flower self-pollination increased, both the import and export of outcrossed pollen in straight-styled plants declined and this shifted the advantage in favour of enantiostylous systems. These results emphasize the importance of herkogamy as a mechanism that reduces forms of self-interference causing self-pollination, particularly in species lacking genetic self-incompatibility (Webb and Lloyd, 1986; Barrett, 2002a). However, it is important to note that under pollen-limited conditions, straight-styled plants may retain an advantage if they are at least partially self-compatible. When outcross pollen receipt is insufficient, self-pollen can offer reproductive assurance as long as inbreeding depression is not severe (Lloyd, 1992; Kalisz et al., 2004; Eckert et al., 2006; Busch and Delph, 2012). Straight-styled morphologies are the dominant stylar condition in angiosperms occurring in virtually all angiosperm lineages and environments. In contrast, enantiostylous species are geographically restricted by comparison.

Our results further suggest that outcrossed pollen receipt is higher in enantiostylous plants compared to straight-styled plants, with the dimorphic condition outperforming the monomorphic condition. (Figure 3). This finding highlights the functional importance of floral asymmetry coupled with reproductive organ reciprocity in promoting disassortative pollination, a finding supported by both theoretical models (Jesson et al., 2003a; Saltini et al., 2024) and empirical studies (Jesson and Barrett, 2002a; Minnaar and Anderson, 2021). In enantiostylous species that possess self-incompatibility systems such as *W. paniculata* (Ornduff and Dulberger, 1978; S. McCarren unpubl. data) this increased capacity to receive outcrossed pollen could directly translate into higher seed set, especially under conditions in which pollinator activity does not limit fertility.

As predicted, the advantage of monomorphic enantiostyly over straight-styled conditions in self-incompatible plants was most pronounced under conditions where within-flower self-pollination is expected to be especially costly, namely when pollinators visit few flowers per plant or deposit only small amounts of pollen (Karron et al., 2003). Conversely, this advantage diminishes under conditions where the relative costs of geitonogamy become greater, such as when pollinators visit many flowers per plant or deposit a larger fraction of available pollen. This pattern supports earlier suggestions that the adaptive significance of enantiostyly depends strongly on the pollination environment (Jesson and Barrett, 2002b). Under these latter conditions in which geitonogamous self-pollination increases, dimorphic enantiostyly may be particularly advantageous. By promoting reciprocal pollen placement, dimorphism not only reduces geitonogamous selfing but also enhances cross-pollen transfer and pollen carryover, allowing this form of stylar polymorphism to maintain a reproductive advantage across a broad range of pollination environments (Barrett, 2002b).

The much more common monomorphic enantiostyly may represent a strategy that alleviates within-flower selfing when displays are small, whereas dimorphic enantiostyly provides a stronger safeguard against the cumulative costs of both within-flower selfing and geitonogamy when floral display size or pollinator visitation rates increase. The reproductive flexibility of dimorphic enantiostyly adds to the puzzle discussed earlier on why this reproductive polymorphism is so rare in angiosperms, given its apparent ecological and fitness benefits. One possibility aside from developmental and selective constraints is that the high levels of disassortative mating that dimorphic enantiostyly promotes essentially reduces the number of potential mating partners in a population by half compared with monomorphic enantiostyly. Such a constraint may limit genetic diversity because of the limit on admixture and under some conditions this may limit opportunities for local adaptation not faced by populations of monomorphic enantiostyly where in principle all individuals in a population are potential mating partners.

From a macro-evolutionary perspective, these results suggest that monomorphic and dimorphic enantiostyly might represent sequential stages along an adaptive pathway shaped by pollination ecology (Fig. 6). Monomorphic enantiostyly could arise first in lineages with small floral displays, where the primary challenge is limiting within-flower selfing (Dulberger and Ornduff, 1980; Harrison et al., 1999). Once floral display sizes increase or pollinator visitation intensifies, selection may then favour the evolution of dimorphic enantiostyly as a more effective mechanism for reducing geitonogamy and maximizing pollen transfer efficiency (Jesson and Barrett, 2002b). This stepwise scenario aligns with comparative evidence that dimorphic enantiostyly has evolved from monomorphic states (Barrett, 2002b; Jesson and Barrett, 2003) and highlights how shifts in floral display and pollination environment have the potential to drive evolutionary transitions in floral symmetry.

**Figure 6.**
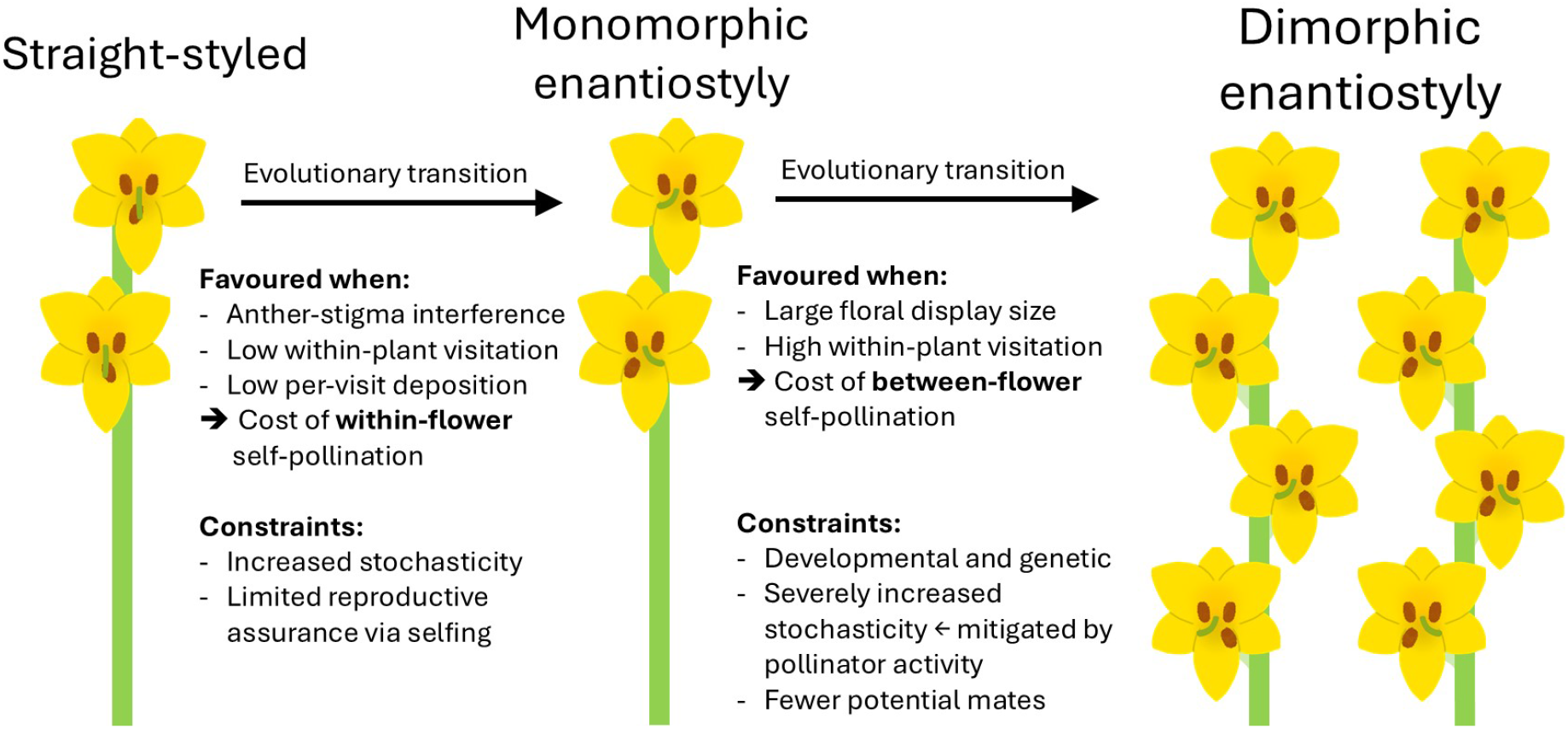
Evolutionary pathways of enantiostyly: selective conditions favouring transitions from straight styles to monomorphic enantiostyly and secondary transitions from monomorphic to dimorphic forms.

Finally, we determined that the optimal fraction of pollen deposited on a focal stigma to maximize cross-pollination differed between the stylar conditions in our study (Fig. 3B). For straight-styled plants, the optimal deposition rate for maximum outcrossing was approximately 20 %, whereas for enantiostylous plants it was closer to 2 %. These findings are intriguing in light of the floral morphology of many enantiostylous species which have minute stigmatic surfaces, which might reflect an adaptation reducing stylar clogging and promoting more precise cross-pollination (Barrett et al., 2000). By limiting the surface area for pollen deposition, enantiostylous flowers may fine-tune pollen receipt in a way that favours disassortative pollination as well as discouraging both within-flower and geitonogamous self-pollination, even when pollinators visit many flowers per plant.

The results of this study support the idea that enantiostyly, particularly dimorphic enantiostyly, enhances fitness mediated through both pollen receipt and export by increasing the precision and distance of outcross pollen transfer while at the same time reducing the cost of within-flower and geitonogamous self-pollination. Although these reproductive benefits may come at the cost of increased vulnerability to variable pollination environments, overall gains in siring success and mate diversity clearly make it an evolutionarily stable strategy in certain ecological contexts and in angiosperm lineages capable of evolving left-right asymmetries in floral structures.

Our mathematical model complements recent theoretical models in pollen transfer (e.g., Saltini et al., 2025; Revilla, 2025; Morita et al., 2025; Mailly et al., 2025) by accounting for a broad repertoire of pollen movements, including pollen import, export, geitonogamous transfer, and intrafloral self-pollination. At the same time, we have not addressed interspecific pollen transfer, which can result in reduced fitness (Revilla, 2025; Morita et al., 2025), nor do we model the co-dynamics of pollen and nectar that may jointly influence pollinator decisions (Revilla et al., 2025). Likewise, our models assume random pollinator movement rather than incorporating reinforcement learning and adaptive foraging behaviour as explored in the agent-based model by Mailly et al. (2025). By integrating diverse forms of pollen transfer within a unified framework, our model provides a foundation that can be extended to include these additional processes. In doing so, it helps to bridge the gap between simple carryover formulations and more mechanistic, behaviorally explicit models of the pollination process.

## Acknowledgements

Permission for this work was granted by Cape Nature permit no. CN35-87-25844.

## Supplementary Figures

**Figure S1.**
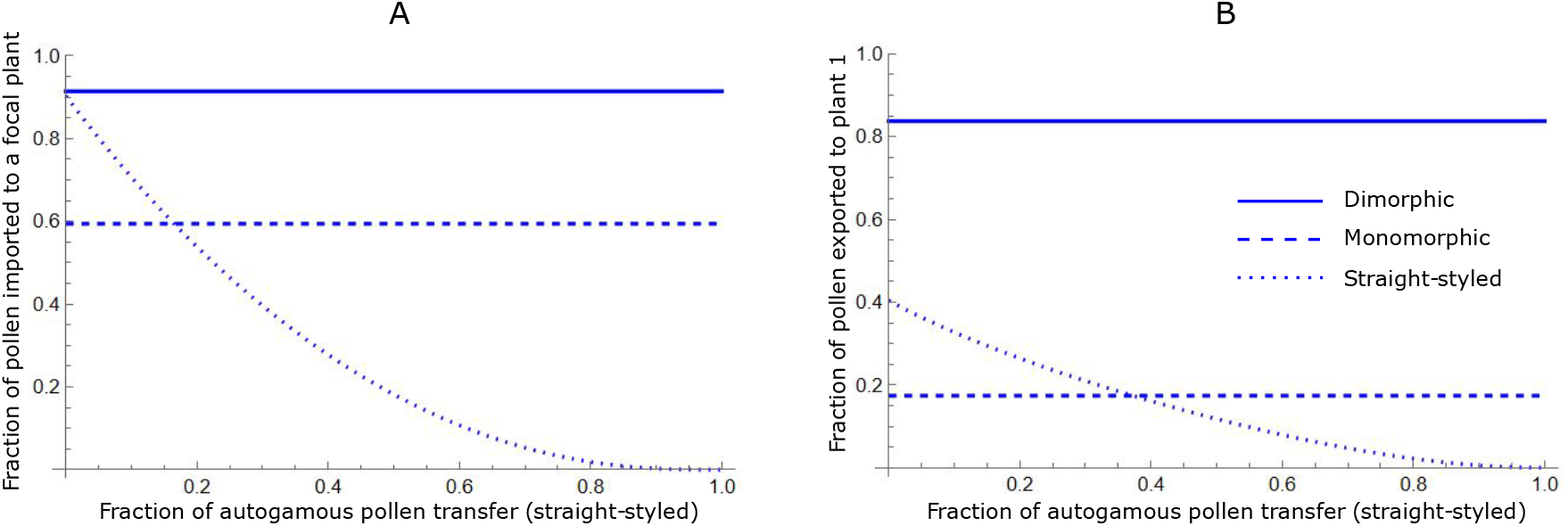
Model results of the average fraction of outcrossed pollen (A) received by a focal plant and (B) exported to the subsequent plant in the pathway of a pollinator, as functions of the fraction of intrafloral pollen transfer in straight-styled species (*λ*_*S*_). Solid lines represent dimorphic enantiostyly, dashed lines represent monomorphic enantiostyly, and dotted lines represent straight-styled species.

**Figure S2.**
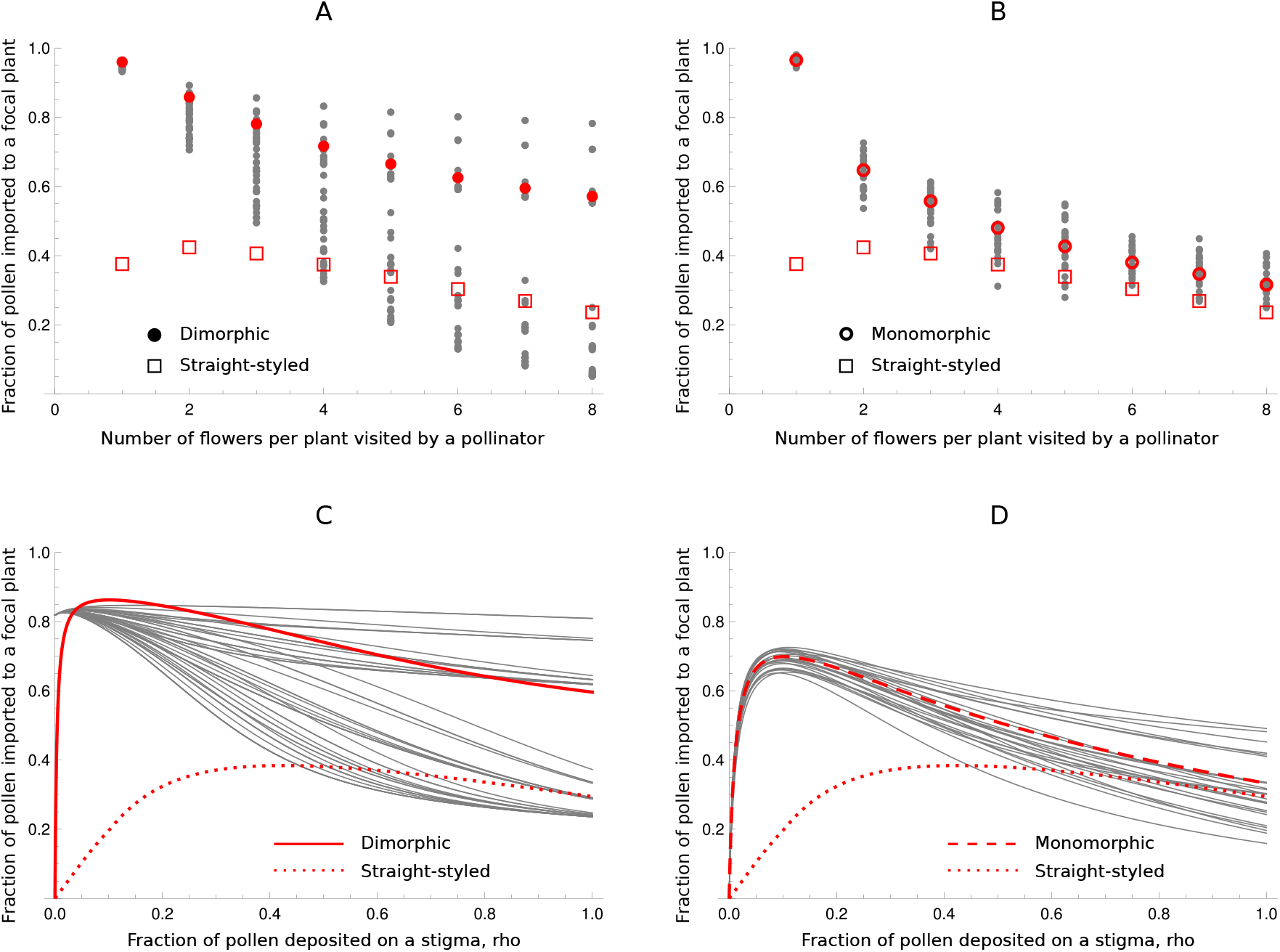
Model results of the average fraction of outcrossed pollen received by a focal plant (red large circles and squares, and red thick lines) calculated over all possible pollinator pathways as a function of: (A, B) the number of flowers visited per plant by the pollinator (*v*), and (C, D) the fraction of pollen on the body of a pollinator deposited on the stigma of the focal plant (*ρ*). Solid lines and filled circles represent dimorphic enantiostyly, dashed lines and empty circles represent monomorphic enantiostyly, and dotted lines and empty squares represent straight-styled species. Grey circles and lines represent fraction of pollen import on single pathways in the (A, C) dimorphic and (B, D) monomorphic cases.

